# Observing action sequences elicits sequence-specific neural representations in frontoparietal brain regions

**DOI:** 10.1101/356253

**Authors:** Dace Apšvalka, Emily S. Cross, Richard Ramsey

## Abstract

Learning new skills by watching others is important for social and motor development throughout the lifespan. Prior research has suggested that observational learning shares common substrates with physical practice at both cognitive and brain levels. In addition, neuroimaging studies have used multivariate analysis techniques to understand neural representations in a variety of domains including vision, audition, memory and action, but few studies have investigated neural plasticity in representational space. As such, although movement sequences can be learned by observing other people’s actions, a largely unanswered question in neuroscience is how experience shapes the representational space of neural systems. Here we combined pre- and post-training fMRI sessions with six days of observational practice to examine whether the observation of action sequences elicits sequence-specific representations in frontoparietal brain regions and the extent to which these representations become more pronounced with observational practice. Our results showed that observed action sequences are modelled by distinct patterns of activity in frontoparietal cortex and that such representations largely generalise to very similar, but untrained, sequences. These findings advance our understanding of what is modelled during observational learning (sequence-specific information), as well as how it is modelled (reorganisation of frontoparietal cortex is similar manner to that of physical practice). Thus, on a more fine-grained neural level than demonstrated previously, we show the representational structure of how frontoparietal cortex maps visual information onto motor circuits to order to enhance motor performance.

**Significance statement:** Learning by watching others is a cornerstone in the development of expertise and skilled behaviour. However, it remains unclear how visual signals are mapped onto motor circuits for such learning to occur. Here we show that observed action sequences are modelled by distinct patterns of activity in frontoparietal cortex and that such representations largely generalise to very similar, but untrained, sequences. These findings advance our understanding of what is modelled during observational learning (sequence-specific information), as well as how it is modelled (reorganisation of frontoparietal cortex is similar manner to that of physical practice). More generally, these findings demonstrate how motor circuit involvement in the perception of action sequences shows high fidelity to the physical performance of action sequences.

## Introduction

From learning to use chopsticks to dancing the lead role in Swan Lake, humans display a remarkable ability to learn complex new motor skills by watching others perform these actions. However, it remains unclear how visual signals are mapped onto motor circuits for such learning to occur. Indeed, our understanding of how action representations develop during motor learning through physical compared to observational practice remains in its infancy (Frey & Gerry, 2006; Hodges et al., 2007; Ostry & Gribble, 2016; McGregor et al., 2016; Vogt et al., 2007). Here we advance understanding of observational learning by using functional magnetic resonance imaging (fMRI) to test the idea that observational learning of action sequences leads to distinctive patterns of activity in sensorimotor cortices, in a manner similar to that reported following physical practice (Wiestler & Diedrichsen, 2013).

Common brain regions have been shown to underpin motor learning following physical and observational experience (Cross et al., 2009; Kirsch & Cross, 2015; Ostry & Gribble, 2016). For example, if the motor system is engaged in another task (Mattar and Gribble, 2005) or if sensorimotor systems are disrupted through non-invasive stimulation (Brown et al., 2009; McGregor et al., 2016), observational learning is reduced. Further, neuroimaging studies have demonstrated that frontoparietal cortex shows similar changes in magnitude and connectivity when learning through physical and observational practice (Cross et al., 2009; Vogt et al., 2007; Higuchi et al., 2012; Sakreida et al., 2018; van der Helden et al., 2010). While these studies demonstrate that sensorimotor cortices are involved in learning motor skills by observation, it remains unclear *how* visual signals are mapped onto motor circuits for learning to occur.

Compared to action observation and visual training, considerably more research has investigated neural representations underpinning action execution and physical training (Dayan & Cohen, 2011; Diedrichsen & Kornysheva, 2015; Hardwick et al., 2013; Kelly & Garavan, 2005; Penhune & Steele, 2012). fMRI studies have shown both increases and decreases in frontoparietal cortex following motor learning, with increases argued to reflect recruitment of cortical tissue and decreases suggestive of more efficient neural function (Dayan & Cohen, 2011; Gardner, Aglinskas & Cross, 2017; Steele and Penhune, 2010). However, because conventional fMRI analyses average activity across voxels, they are insensitive to a richness of information that is represented by the pattern of activity across voxels (Kriegeskorte et al., 2008). Sidestepping the issue of averaging across voxels, Wiestler and Diedrichsen (2013) used a motor learning paradigm in combination with multi-voxel pattern analysis (MVPA) to identify how patterns of activity across voxels relate to mental content, independent of average activity (Norman et al., 2006; Kriegeskorte et al., 2008). Wiestler and Diedrichsen (2013) showed that execution of kinematically-matched keypress sequences was associated with sequence-specific patterns of activity in multiple frontoparietal brain areas. Moreover, physically practicing sequences led to reduced activity on average and more distinctive patterns of activity in frontoparietal brain areas, implying a more distinct neural representation of learned sequences that enables faster execution (Wiestler and Diedrichsen, 2013).

To date, MVPA has been used to understand neural representations in a variety of domains including vision, audition, memory and action, but few studies have investigated neural plasticity in representational space (Kriegeskorte & Kievit, 2013). Therefore, although movement sequences can be learned by observing other people’s actions (Bird et al., 2005; Blandin et al., 1999; Boutin et al., 2010; Hodges et al., 2007; Vogt et al., 2007), a largely unanswered question in neuroscience is how experience shapes the representational space of neural systems (Kriegeskorte & Kievit, 2013). To address this question, here we test the extent to which observation of action sequences elicits sequence-specific representations in frontoparietal brain regions and the extent to which these representations become more pronounced with observational practice. If observed sequences are mapped onto sensorimotor circuits in a similar manner to physical practice (Wiestler and Diedrichsen, 2013), we would expect sequence-specific patterns of activity to emerge within sensorimotor cortices following observational training.

## Method

### Participants

Eighteen right-handed (based on self-report) volunteers from the Bangor University student community participated in the study. Two participants were not included in the final sample: a pilot participant, who did not have the same testing parameters, and a participant who made excessive head movements during one of the scanning sessions (> 4 mm). The final sample comprised 16 participants (8 males and 8 females), 20 to 40 years old (M = 24.31 years, SD = 5.06). All participants had normal or corrected-to-normal vision and no history of neurological disorders. Participants gave their written informed consent and were paid £45 for their participation. All procedures were approved by the Ethics Committee of the School of Psychology at Bangor University and UK Ministry of Defence Research Ethics Committee.

### Stimuli

A keypress sequence learning paradigm was implemented, based on the task used by Wiestler and Diedrichsen (2013). A standard QWERTY black computer keyboard was used with the Q 3 4 5 and Y keys covered with red tape and all surrounding keys removed. In pre- and post-training sessions, participants were required to press the red keys with the five fingers of their left hand in a specified order. During the observational training and fMRI sessions, participants watched videos of the experimenter performing the keypress task. For the video recordings, a similar keyboard was used with the only difference that the sides of the five keys were covered in yellow to improve the visibility of the key being pressed. Stimuli presentation and response recordings were performed using MATLAB 8.3.0 (The MathWorks, MA, USA) and Psychophysics Toolbox 3.0.12 (Brainard, 1997).

#### Keypress sequences

The same set of 12 five-element keypress sequences was used as previously by Wiestler and Diedrichsen (2013). Each sequence required the five fingers of the left hand to be pressed once in a sequential order, with each of the 12 sequences featuring a different order with no more than three adjacent finger-presses in a row. All sequences were matched for difficulty, based on a pilot experiment (Wiestler and Diedrichsen, 2013). For each participant, from the set of 12 sequences, four sequences were randomly allocated to the Trained condition, and four other sequences were allocated to the Untrained condition. The remaining four sequences remained used.

#### Videos

For observational training and both scanning sessions, 13-second videos were created showing the experimenter’s left hand from a first-person perspective, slightly tilted to the right (Figure 1C; see Stimuli, https://osf.io/jz4nk/). Each video showed the experimenter executing one sequence five times, with naturally varying breaks between each sequence repetition, to ensure a more authentic presentation of the performance. For the same reason, for each sequence, five different video versions were recorded, to allow closer to natural performance variation of the same sequence. An additional video version for each sequence was created where one of the five sequence executions was incorrect. This resulted in 72 videos in total.

**Figure 1.**
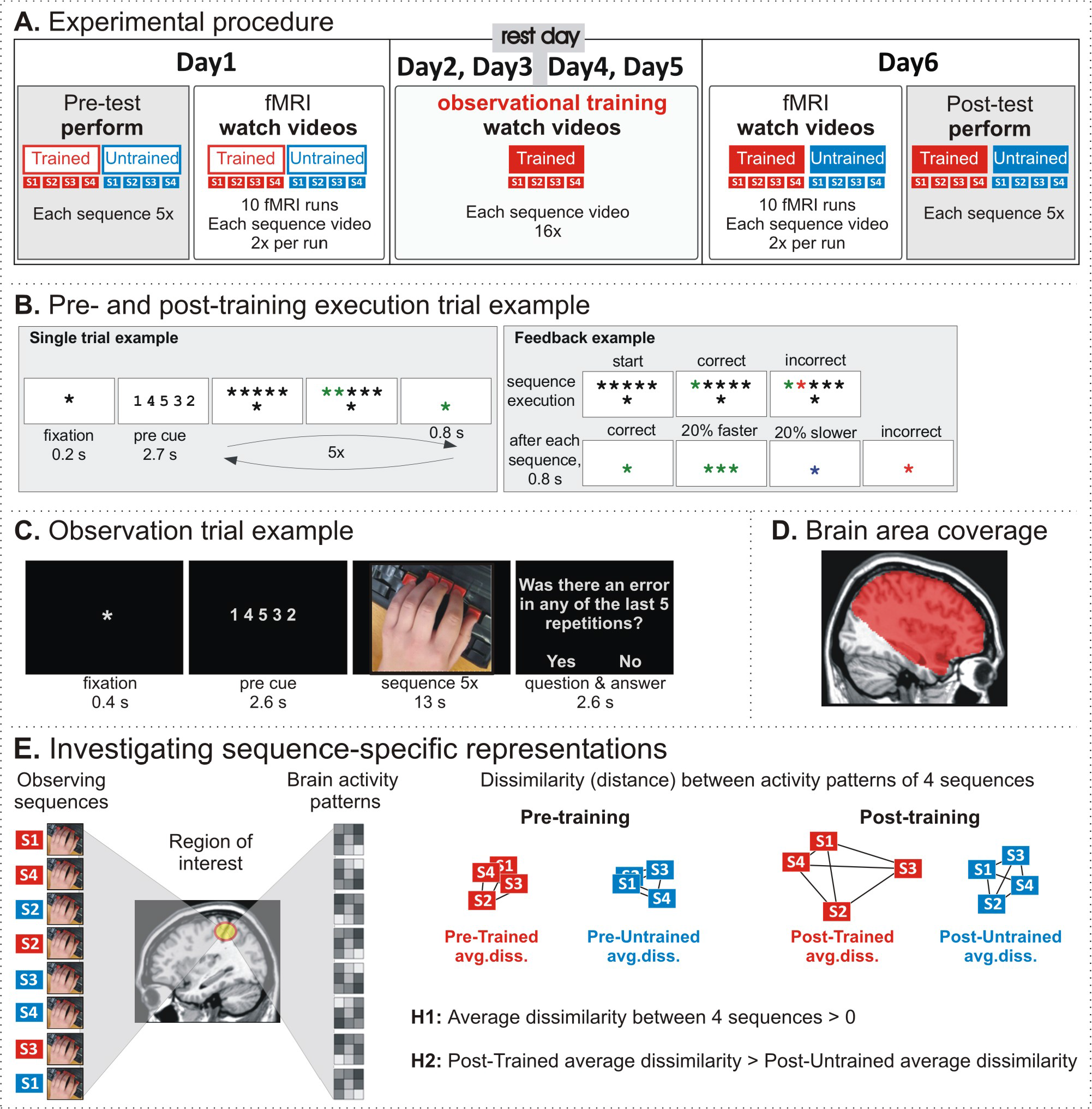
Experimental paradigm. **A.** Experimental procedure. The experiment involved pre-test and post-test, separated by four training days and two scanning (fMRI) sessions. In the pre- and post-test, participants performed eight keypress sequences (four of them to be trained, the other four untrained). In the scanning sessions, participants watched videos of a hand performing the same eight sequences. In the training sessions, participants watched videos of a hand performing four of the eight sequences. **B.** Execution trial example. A cued sequence had to be memorised and then executed five times while receiving performance feedback. **C.** Observation trial example. A sequence cue was followed by a video showing a hand executing the sequence five times, either correctly or incorrectly. Occasionally a question was asked whether there was an error in any of the five repetitions, and a response had to be made. **D.** Brain area coverage for fMRI analysis focused on premotor and parietal brain regions, and did not include the cerebellum, occipital lobes, or inferior temporal lobes. **E.** During pre- and post-training fMRI sessions, participants watched videos of 8 different sequence executions; 4 sequences belonged to Trained and 4 others to Untrained conditions. Within each condition and each fMRI session, we measured the dissimilarity between each pair of the 4 sequences (six pairs) and obtained the average dissimilarity estimate (linear discriminant contrast; LDC) between the 4 sequences. The dissimilarity measures were used to investigate our main hypotheses: 1) action observation evokes movement-sequence-specific brain activity patterns (the average dissimilarity between 4 sequences is above zero); 2) the activity patterns become more distinct following observational practice (the average dissimilarity between 4 trained sequences is higher than between 4 untrained sequences).

Sequences were executed at an intermediate performance level, which was determined by behavioural pilot test results, where the average time to complete a correct sequence execution was 2.29 seconds (N = 17, M = 2.29 s, SE = 0.14). Each original video, showing five repetitions of the same sequence, was slightly speeded up or slowed down (±10%) to make it exactly 13 seconds long. Consequently, the authenticity of movement performance was somewhat reduced, but the relative variability within the video remained intact. The average single sequence execution in the videos was 2.3 seconds. The videos were presented on a computer monitor in full colour on a black background. The frame rate was 29 frames per second with the resolution of 600 × 526 pixels, showing approximately natural hand size.

### Procedure

#### Overview

Participants underwent six testing days over a seven-day period (Figure 1A). On the first day of testing, participants received task instructions and completed three single-sequence execution trials to ensure that each participant understood the task. The familiarisation procedure was followed by a pre-training session, which was immediately followed by the first scanning session. Participants returned to the lab for the next two consecutive days for observational training sessions, which were followed by a day off. After the rest day, participants returned to the lab for two more consecutive days of observational training sessions. The final day (Day 6) started with the second scanning session, immediately followed by a post-training session. Each session is described in more detail below.

#### Pre- and post-training sessions

In the pre- and post-training sessions, participants performed four Trained and four Untrained sequence execution trials in a random order with their left hand. Each trial consisted of five repetitions of the same sequence (Figure 1B). All trial-related information was presented centrally at the bottom of the screen against a grey background. A trial started with a black fixation cross (0.2 s), followed by the sequence cue presented as five digits (2.7 s) that indicated from right to left which key to press: “1” – the right-most key pressed with the thumb; “5” – the left-most key pressed with the little finger. After the cue, the digits were replaced by the fixation cross and five black asterisks above it. This served as a “go” signal to execute the memorised sequence five times as quickly and accurately as possible. If the correct key was pressed, the corresponding asterisk on the screen turned green, if a wrong key was pressed, the asterisk turned red.

After executing a single sequence, the central fixation cross changed colour to provide feedback on the performance (0.8 s): green – correct sequence execution; red – incorrect sequence execution; blue – correct, but executed 20% slower than the median execution time in the previous trials; three green asterisks – correct and executed 20% faster than the median execution time in the previous trials. After this short feedback, all asterisks turned black signalling the start of the next execution trial. After five executions of the same sequence, the trial ended and the next sequence was cued.

#### Observational training sessions

In the observational training sessions, participants watched videos of the four Trained sequence executions. Participants were instructed to watch the videos and to pay close attention to whether the sequences were performed correctly. Occasionally they would be asked whether the performer in the video made an error in any of the five repetitions – the error question. They would respond by pressing the ‘b’ key (marked red) on a keyboard for yes and the ‘m’ key (marked blue) for no. This task was included to ensure that participants paid close attention to the videos. Participants were also informed that they will need to perform the watched sequences again at the end of the experiment.

All trial-related information was presented in the middle of the screen against a black background with a light grey font (Figure 1C). A trial started with a fixation cross (0.4 s), followed by the sequence cue presented as five digits (2.6 s), followed by the sequence video (13 s). After some of the trials, the error question was asked and participants had 2.6 seconds to respond.

A training session was divided into four blocks, separated by a rest period. Within each block, 20 videos were presented in a random order. Each of the four training sequence videos was shown four times (randomly choosing one of the five video versions for each sequence, described in the Videos section above). There was also one ‘error video’ for each sequence (where at least one of the five repetitions of the sequence execution was incorrect). The error question appeared randomly 5-7 times per block. At the end of each block, participants received feedback on how accurately they spotted the incorrect sequence executions. The whole training session lasted approximately 25 minutes and participants saw a correct execution of each sequence at least 80 times (4 blocks, 4 distinct sequence videos per block, 5 repetitions of a single sequence per video, plus some correct repetitions in the ‘error video’).

#### Scanning sessions

During identical pre- (Day 1) and post-training (Day 6) fMRI sessions, participants observed the four Trained and four Untrained sequence videos in a random order. The observation trials were structured in the same way as in the observational training sessions (Figure 1C). In each scanning session, participants completed 10 functional runs. Each functional run comprised 17 videos presented in a random order: eight sequence videos presented twice each, and one ‘error video’. Each video showed five repetitions of one sequence. Therefore, during each scanning session, participants saw a correct execution of each sequence at least 100 times (10 functional runs, 2 videos per sequence per run, 5 repetitions of a single sequence per video, plus some correct repetitions in the ‘error video’).

In keeping with the observational training sessions, participants were instructed to watch whether all sequences were correctly executed and to answer the error question when asked. The error question was asked twice within a run – always after the ‘error video’ and randomly after one of the correct videos. Each run also had five rest phases, one at the beginning of the run and four randomly interspersed, but never twice in a row. The rest phase was 13 seconds long and showed a fixation cross in the middle of the screen. Each run lasted approximately 6 minutes (2.6 s per whole-brain acquisition, with 138 acquisitions per run).

Stimuli were presented onto a screen located behind the MRI scanner and displayed to the participant via a mirror positioned above participants’ eyes. Responses to the error questions were recorded using a scanner-safe fibre optic four-button response pad (Current Designs, Philadelphia, PA) connected to the stimulus PC.

### Scan acquisition

MRI data were acquired using a 3 Tesla Phillips Achieva MRI scanner (Philips Health Care, Eindhoven, Netherlands) fitted with a sensitivity-encoded (SENSE) 32-channel phased-array head coil.

#### Functional scans

Both scanning sessions consisted of 10 functional runs of the blood-oxygenation-level-dependent (BOLD) signal acquisitions (Ogawa et al., 1992), with two dummy scans and 136 whole-brain scans per run. Volumes were collected using a T2*-weighted single shot gradient echo planar imaging sequence with the following parameters: TE = 30 ms, TR = 2.6 s, flip angle = 90°, 41 ascending slices with 2.3 mm thickness, 0.15 mm gap, and 2 × 2 mm^2^ in-plane resolution (matrix size 96 × 96). The slice acquisition was focused on premotor and parietal brain regions, thus the group average brain area coverage did not include the cerebellum, or all of the occipital or inferior temporal lobes (Figure 1D).

#### Anatomical scan

The last scanning session (Day 6) ended with a high-resolution whole-brain 3D anatomical scan acquired as a T1-weighted image (MP-RAGE, TE = 3.5 ms, TR = 12 ms, voxel resolution = 1 mm^3^, slice thickness = 2 mm, flip angle=8°), which was used as an anatomical reference for each participant.

### Data analysis

#### Overview of analysis strategy

The general analysis strategy is motivated by our main research question, which is focussed on understanding how changes in the pattern of activity in frontoparietal cortex supports sequence-specific representations following observational learning. More specifically, we measure the extent to which individual observed action sequences are represented by patterns of activity in frontoparietal cortex, as well as the extent that these sequence-specific representations are dissociable in a training-specific manner (i.e., trained > untrained). In addition, we focussed our pattern classification analyses on specific region of interest (ROIs) that, as measured by average activity across voxels, showed sensitivity to observational learning during the post-training scan session (e.g. Sakreida et al., 2018). By focussing our pattern analyses on regions that satisfy functional criteria associated with observational learning, we ensure that inferences drawn regarding representational-level and sequence-specific effects are in brain regions that are sensitive to observational learning.

The analyses performed within these ROIs closely follows analyses reported in prior physical training studies that have employed a sequence learning task (Wiestler & Diedrichsen, 2013; Wiestler et al., 2014). In terms of behavioural effects following sequence learning, Wiestler and colleagues (2013; 2014) reported skill learning that generalised across all sequences (significant pre- to post-training performance improvement of both trained and untrained sequences) and training-specific sequence learning (greater post-training performance for trained than untrained sequences).

In addition, in frontoparietal brain regions, measures of average activity and MVPA showed evidence for generalised skill learning and training-specific effects (Wiestler & Diedrichsen, 2013). Here, we performed similar analyses of behavioural and brain data to test the extent to which observational training effects generalise across trained and untrained sequences, and whether these effects dissociate between trained compared to untrained sequences. To do so, we first assessed sequence-specific learning for trained and untrained sequences separately. That is, we assessed the extent to which distinctive patterns of activity for observed action sequences (irrespective of training condition) are identifiable within task-defined regions of the frontoparietal cortex. This first analysis is an important extension to prior sequence-learning action observation studies that used univariate measures (e.g., Frey & Gerry, 2006; Sakreida et al., 2018), as univariate measures are unable to distinguish between the neural representation of individual sequences. Indeed, univariate measures can distinguish between a collection of trained and untrained sequences, but the coarseness of univariate measures does not allow individual sequences to be distinguished.

Second, to test the extent to which sequence-specific patterns of activity in frontoparietal cortex dissociate between sequences in a training-specific manner, we assessed differences between representations of trained and untrained sequences during the post-training scan session. To correct for possible pre-training differences, we followed the approach by Wiestler and Diedrichsen (2013) and calculated a linear regression between the pre-training difference (predictor) and the posttraining difference (outcome). The intercept of the regression line was used as a measure of the post-training difference between Trained and Untrained conditions, correcting for possible pretraining differences. The linear regression approach was used in all subsequent behavioural and brain imaging analyses (univariate and MVPA) when comparing Trained and Untrained conditions after training.

Finally, to complement these ROI-based pattern analyses, we also performed a whole-brain searchlight analysis. Because sequence-specific representations of observed action sequences have not been investigated before, the whole-brain searchlight analysis enables us to characterise our main research questions beyond our ROIs.

#### Behavioural performance

Participants’ physical performance was assessed pre- and post-training, measuring the average sequence initiation time, execution time and error rate of the four trained (to-be-trained) and the four untrained sequences. The sequence initiation time was measured as the duration between the “go” signal and the first keypress. The sequence execution time was measured as the duration between the first and fifth keypresses. The error rate was measured as the percentage of incorrect sequence executions. Incorrectly executed trials were excluded from further analysis. Attention to the task during the observational training and scanning sessions was assessed as a percentage of accurate responses to questions on error trials.

#### Imaging data

Imaging data were analysed using SPM12 (Wellcome Trust Centre for Neuroimaging, London), and custom-written MATLAB scripts. To correct for head motion, all images from a single scanning session (10 × 136 volumes) were spatially realigned to the mean functional image and slice-time corrected. The anatomical T1-wighted image was co-registered to the session-mean functional image and segmented to obtain parameters for spatial normalisation. The time series of each voxel were high-pass filtered with a cut-off frequency of 1/52 Hz, to remove low frequency trends, and modelled for temporal autocorrelation across scans with an AR(1) process.

For the voxel-wise univariate analysis, normalisation parameters from the segmentation step were used to normalise pre-processed functional images to the Montreal Neurological Institute (MNI) template brain with a resolution of 2 mm^3^. Normalised images were then spatially smoothed with a 3D Gaussian kernel of 8 mm full-width-half-maximum (FWHM). MVPA was performed without normalisation and smoothing, to preserve high spatial resolution.

All statistical maps were thresholded at a single voxel level with a significance value of p <0.001 and a minimum cluster size of 10 voxels. To control for false positive results, only brain regions reaching cluster familywise error (FWE) corrected significance at p < 0.05 are reported. For anatomical and cytoarchitectonic localisation, we used SPM Anatomy toolbox v2.0 (Eickhoff et al., 2005).

#### Univariate analysis

The univariate analyses were designed to achieve two main objectives: (1) identify brain regions engaged in action observation, and (2) identify brain regions sensitive to observational practice. Normalised and smoothed data were analysed using a General Linear Model (GLM). A random-effects model was implemented at two levels. At the first level, single participant data were modelled by a single design matrix for all runs within each session. The design matrix contained 6 regressors of the following events: Trained videos, Untrained videos, an ‘error’ video, error questions/responses, Trained cues, and Untrained cues. Trained and Untrained video regressors (further named, Trained and Untrained) represented the 13-second video duration (showing five repetitions of a single sequence execution). All regressors were modelled as boxcar functions, convolved with a hemodynamic response function (HRF). The rest periods formed an implicit baseline.

To identify brain regions engaged in action observation, only data from the pre-training scanning session were used. Both action observation conditions of the pre-training session were taken together and contrasted with the implicit baseline (pre-Trained ⋃ pre-Untrained > implicit baseline). The first level whole-brain contrast maps where then entered into a second-level one-sample t-test analysis to obtain group average results of brain areas engaged when watching keypress sequences in general, pre-training.

To identify brain regions sensitive to observational practice, the linear regression approach was used, as described above in the ‘Overview of the analysis strategy’ section. Specifically, the pre-training difference between the estimated beta weights of the Trained and Untrained conditions within each of the 10 pre-training functional runs was used as a predictor variable. The post-training difference between the Trained and Untrained conditions within each of the 10 post-training functional runs was used as an outcome variable. The intercept of the regression line was used as a measure of the post-training difference between the Trained and Untrained conditions, correcting for possible pre-training differences. The linear regression was performed at the first level, in a voxel-wise manner across the whole brain and produced the intercept maps for each subject. These first level whole-brain maps where then entered into a second-level one-sample t-test analysis to obtain group average results of brain areas sensitive to observational practice.

#### Region of interest definition

Based on univariate data, peak voxels from significant clusters showing the post-training difference between Trained and Untrained conditions (independent of the direction) were used to create regions of interest (ROIs) for MVPA. We note that our analysis approach is not circular (Kriegeskorte et al., 2009), because the univariate analysis of post-training differences is statistically independent to all subsequent analyses.

More specifically, the ROIs were defined for each participant as follows (Figure 2). First, 15 mm radius spheres centred on the group level voxels with the highest t-value of the post-training difference were created in MNI space (these ROIs are available at http://neurovault.org/collections/1892/). Second, at an individual participant level, voxels with the highest post-training difference value within the 15 mm radius spheres were selected as the individual’s peak voxels. This approach was taken to account for anatomical and functional variability in the areas responsive to the task across participants. Third, 10 mm radius spheres centred on the individuals’ identified peak voxels were created for beta weight extraction to visualise the response. Fourth, the 10 mm radius spheres were mapped from the MNI space onto individual subject anatomies for MVPA analysis.

**Figure 2.**
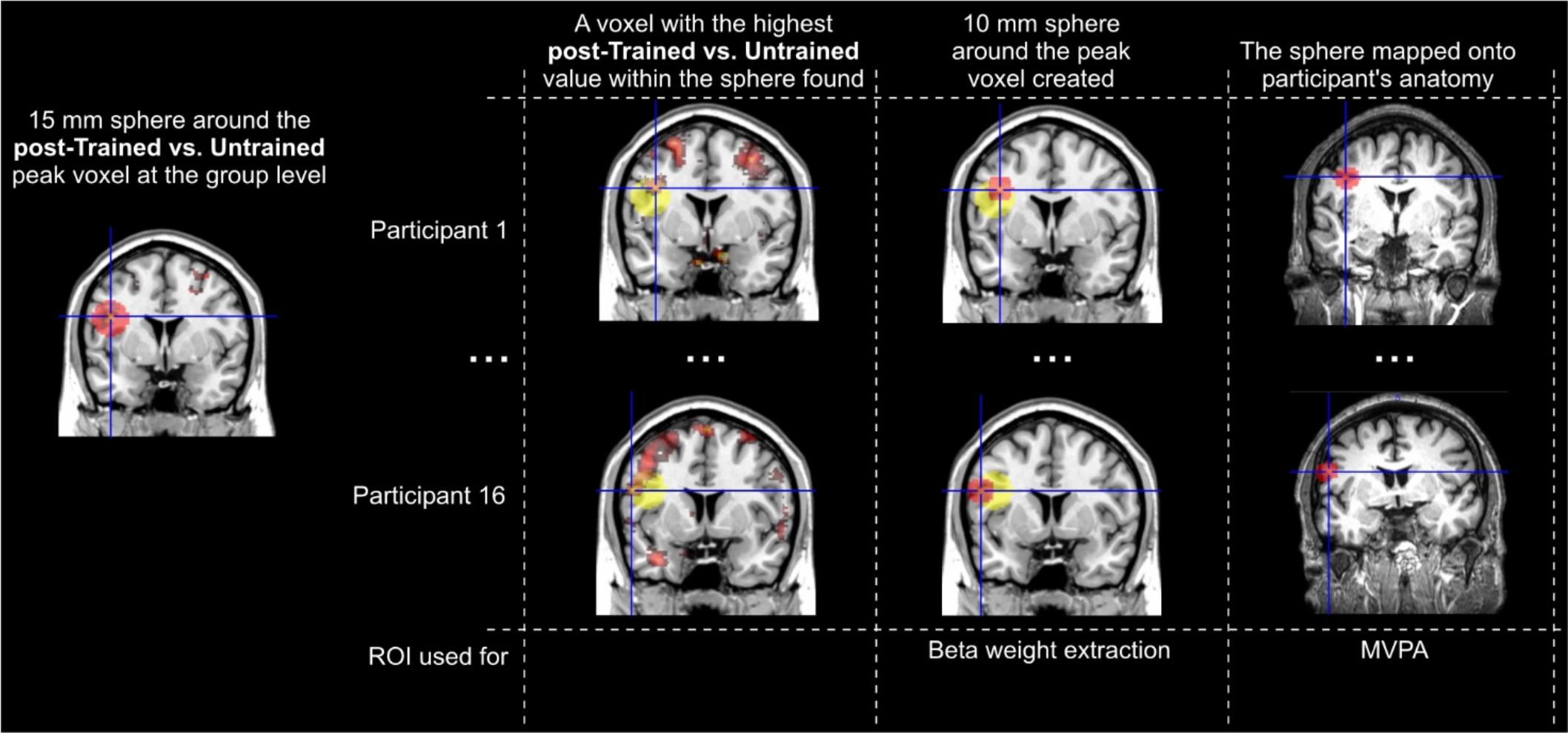
Region of interest (ROI) definition procedure. The peak voxels of significant clusters showing the training-related brain activity changes were selected for ROI based functional connectivity and MVPA analyses. **First,** 15 mm radius spheres were created in the MNI space, centred on the group level voxels with the highest t-value of the posttraining difference between Trained and Untrained conditions (independent of the direction). **Second,** at a participant level, each individual’s peak voxels were identified within the group level 15 mm radius spheres. **Third,** 10 mm radius spheres centred on the identified individuals’ peak voxels were created for beta weight extraction. **Fourth,** the 10 mm radius spheres were mapped from the MNI space onto individual subject anatomies for MVPA analysis.

#### Multi-voxel pattern analysis - region of interest approach

MVPA was implemented to achieve two main objectives: (1) identify brain regions associated with sequence-specific representations through action observation, and (2) identify the extent to which patterns of activity become more sequence-specific following observational training of action sequences. To test whether the observation of action sequences is associated with sequence-specific representations in frontoparietal cortex, we used MVPA to analyse brain activity patterns that emerge when watching the four sequences within each condition (Trained and Untrained). Consistent with the previous physical training study (Wiestler and Diedrichsen, 2013), we first examined sequence-specific patterns within each condition separately to test if neural representations in frontoparietal cortices distinguish between observed key-press sequences in general. Second, we then compared the results across training conditions to determine whether the patterns of activity in frontoparietal cortex become more distinct for trained compared to untrained sequences (Figure 1E).

The dissimilarity between activity patterns was measured using cross-validated Mahalanobis distance (Diedrichsen et al., 2016), which is closely related to linear discriminant analysis (LDA), and therefore termed linear discriminant contrast (LDC). In a recent study, LDC proved to be the most reliable MVPA measure, outperforming other more popular measures, such as pattern classification (LDA and support vector machine) and Pearson correlation (Walther et al., 2016).

LDC is a continuous dissimilarity measure, which includes multivariate noise normalisation (pre-whitening), cross-validation, and does not depend on baseline activity. Similar to LDA, LDC compares two conditions using a linear discriminant that has been estimated with independent data. However, instead of a binary decision, which is then converted into classification accuracy, LDC computes the mean difference between the two conditions measured along the linear discriminant. Cross-validation ensures that the measured dissimilarities between conditions are not due to noise in the data that makes conditions appear to differ by chance, but instead it ensures that dissimilarity measures represent a true difference with a meaningful zero point (Diedrichsen et al., 2016; Walther et al., 2016). If a brain region differentiates between the two types of stimuli (or two conditions), the average cross-validated dissimilarity measure of the activity patterns would be above zero.

Here the LDC analysis was implemented using the RSA Toolbox (Nili et al., 2014) and custom-written MATLAB scripts. To obtain activity patterns for LDC analysis, a first-level GLM was estimated for each participant using the spatially realigned and slice-time corrected images, without normalisation and smoothing. For the pre-training and post-training data separately, a unique regressor for each of the eight sequences (four Trained, four Untrained) within each of the 10 runs was modelled as a boxcar function and convolved with an HRF. Each regressor averaged the brain activity across the two occurrences of the 13-second videos of each sequence within each run.

The LDC analysis of the activity patterns across sequences was performed for each condition (Trained and Untrained) and each participant separately. The estimated beta weights of the voxels in each region (ROI or searchlight) were extracted and pre-whitened to construct noise normalised activity patterns for each sequence within each run (Diedrichsen et al., 2016; Walther et al., 2016). As such, the input data for the LDC analysis consisted of 4 × 10 (four sequences, 10 runs) activation estimates for a set of 160 neighbouring voxels within each ROI. Leave-one-run-out cross-validated LDC analysis was performed, and dissimilarity estimates averaged across the ten possible crossvalidation folds.

For each training condition and ROI separately, we compared patterns of activity between all four observed sequences to each other. This produced a total of six comparisons. For each comparison, we calculated the dissimilarity in patterns of activity as measured by 1 minus the correlation of the activity patterns (Kriegeskorte et al., 2008; Kriegeskorte & Kievit, 2013). Hence, if patterns of activity between two sequences were perfectly correlated, there would be zero dissimilarity. Likewise, lower correlations (similarity) between the patterns of activity between two sequences will produce greater dissimilarity scores. The resulting six dissimilarity scores were averaged to obtain the average dissimilarity estimate between the four sequences. An above-zero dissimilarity estimate indicates that the examined region (ROI or searchlight) has a pattern of activity that represents sequence-specific information.

For MVPA ROI analyses, we used a random subspace approach to increase the reliability of LDC measures (Diedrichsen et al., 2013). To do so, for each ROI separately, subsets of 160 voxels were randomly selected 1000 times. LDC analysis was performed on each subset and dissimilarity estimates from all 1000 subsets were averaged to obtain the final LDC measure for each ROI and each condition: LDC pre-Trained, LDC pre-Untrained, LDC post-Trained, and LDC post-Untrained. Results were then submitted for statistical analyses.

First, we estimated the condition-average sequence-specific coding pre- and post-training separately. To do so, for the pre- and post-training scanning data separately, we averaged the Trained and Untrained LDC values and tested them against zero using one-tailed t-test. An above zero value would indicate that patterns of activity are distinct between sequences. Next, we assessed the post-training difference (intercept) between the training conditions (trained > untrained), correcting for the possible pre-training differences (as described previously). All tests were Bonferroni-corrected for the four ROIs. Accordingly, the significance threshold for the statistical comparisons was p < 0.0125.

#### Multi-voxel pattern analysis – searchlight approach

In an exploratory whole-brain analysis, we performed a surface-based searchlight analysis (Oosterhof et al., 2011) to identify brain regions coding sequence-specific information across the whole cortical surface (Kriegeskorte et al., 2006). Cortical surfaces were reconstructed from individual T1-weighted images using FreeSurfer (Dale et al., 1999). Around each surface node, spheres of searchlights were defined and all voxels between pial and white-grey matter surface selected for analysis. The radius of each sphere was adjusted such that each searchlight contained exactly 160 voxels. The average searchlight radius was 10.37 mm.

For each searchlight, LDC analysis was performed for the four sequences within each condition as described in the MVPA ROI analysis section above. The dissimilarity estimate of each searchlight was assigned to the central voxel, constructing a surface map of dissimilarity estimates. The acquired individual subject maps (LDC pre-Trained, LDC pre-Untrained, LDC post-Trained, and LDC post-Untrained) were then normalised to the MNI template, with a resolution of 2 mm^3^, and spatially smoothed, with a 3D Gaussian kernel of 4 mm FWHM.

The normalised and smoothed maps were then entered into a second-level random-effect analysis to obtain group average results of brain areas that code sequence-specific information when watching sequences pre-training and post-training (one-sample t-tests against zero of LDC preTrained ⋃ LDC pre-Untrained and of LDC post-Trained ⋃ LDC post-Untrained). We also calculated the post-training difference between the Trained and Untrained conditions, correcting for possible pre-training differences, using the linear regression approach as described previously.

In addition, following Wiestler and Diedrichsen (2013) approach, we also inspected the sequence-specific representations globally, averaging over all involved cortical regions.

Specifically, for each participant we created a mask of cortical areas where LDC value was above zero for any of the four conditions. Within this mask, for each condition separately, we calculated the average LDC value and the total area where the LDC value was above zero. Next, individual participant LDC and total area values where entered into the regression analyses to compare the post-training difference between the Trained and Untrained conditions, correcting for possible pretraining differences (as described previously).

### Reported confidence intervals

All sample means are reported with their 95% confidence intervals in square brackets. Confidence intervals for two-tailed tests were calculated as SE * 2.13, whereas confidence intervals for one-sided tests were calculated as SE * 1.74 for df 15 (Cumming, 2012). When reporting results for multiple conditions, within-subject confidence intervals were used (Cousineau, 2005).

### Data sharing

Stimuli, data, and code for this study are freely available at https://osf.io/jz4nk/. In addition, we also performed an exploratory functional connectivity analysis using psychophysiological interactions (PPI) analyses (see PPI_analysis; https://osf.io/jz4nk/). Unthresholded fMRI maps, LDC maps and group ROIs are uploaded at http://neurovault.org/collections/1892/.

## Results

### Behavioural data

We first assessed the extent to which participants were paying attention to the videos during observational training and scanning sessions by analysing accuracy of performance on identifying error videos. The average accuracy across the four training days was 87% [81%, 93%]. On average, accuracy improved across the four training days (Figure 3A), but the difference was not significant, as measured by a 4-way repeated-measures analysis of variance, F_3,42_ = 1.076, p = 0.370. The average accuracy during the scanning sessions was 69% [58%, 80%], with no significant difference between the two sessions, t_15_ = 0.786, p = 0.444, d_z_ = 0.20. Therefore, we can be reasonably confident that participants paid attention to the videos during observational training and scanning sessions.

**Figure 3.**
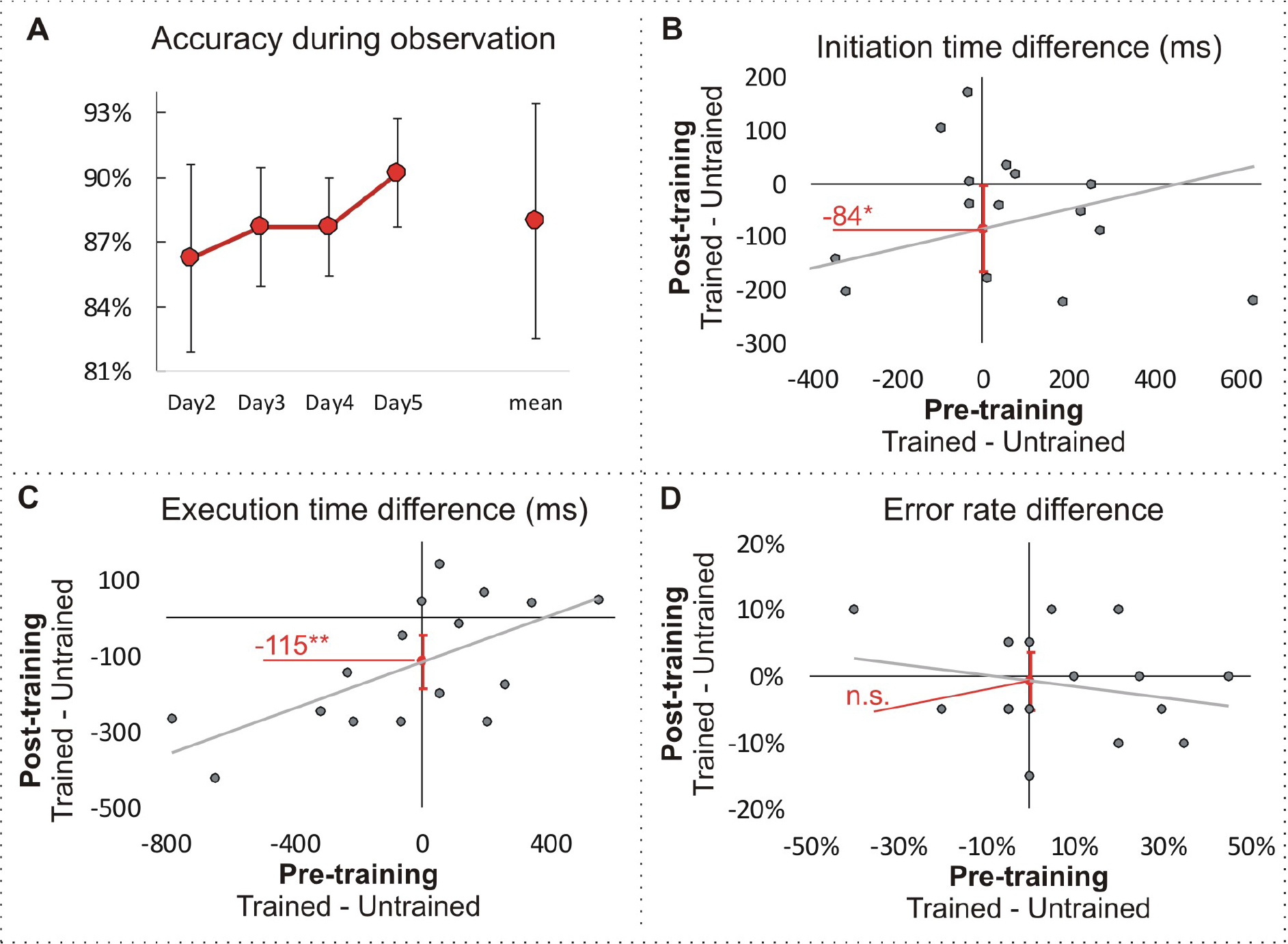
Behavioural results. **A.** Group-averaged accuracy in response to the error question during observational training. Error bars represent within-subject 95% CI.**B.**, **C.** and **D.** Pre- and post-training difference in initiation time, execution time and error rate between trained and untrained sequences. The training effect was measured as the intercept of the regression line between the pre-training difference (predictor) and the post-training difference (outcome). The intercept represents the predicted post-training difference if the pre-training difference is zero. This method reduces the noise of unwanted differences in the difficulty of trained and untrained sequences and thus allows a more accurate measurement of the training effect. Error bars represent 95% CI of the intercept. * p < 0.05, ** p < 0.01, n.s.: non-significant at p < 0.05.

Post-training, sequence initiation time for the trained sequences (M = 600 ms [526 ms, 674 ms]) was significantly faster than for the untrained sequences (M = 684 ms [612, 756]), t_14_ = 2.238, p = 0.042, d_z_ = 0.56, B_0_ = −84 ms [−165, −4] (Figure 3B). Execution time for the trained sequences (M = 1338 ms [1215 ms, 1461 ms]) was significantly faster than for the untrained sequences (M = 1464 ms [1365, 1562]), t_14_ = 3.495, p = 0.004, d_z_ = 0.87, B_0_ = −115 ms [−185, −45] (Figure 3C). Therefore, effects sizes for our primary behavioural measures of observational learning (initiation and execution time) are typically considered medium and large, according to Cohen’s benchmarks (Cohen, 1992). Error rate did not differ between the two conditions (post-Trained M = 12% [7, 18]; post-Untrained M = 13%, [9, 18]), t_14_ = 0.319, p = 0.754, d_z_ = 0.08, B_0_ = −0.6% [−5, 4] (Figure 3D).

### fMRI data

#### Univariate analyses

##### Brain regions engaged in action observation

To identify brain regions engaged when watching sequences in general, a group average contrast of pre-Trained ⋃ pre-Untrained > implicit baseline was assessed. The brain regions that emerged from this contrast included bilateral superior and inferior parietal lobules, intraparietal sulci, dorsal premotor cortices (including supplementary motor area), hippocampi, and left ventral premotor cortex. A list of the major peaks of activated clusters is given in Table 1 and all activated areas visualised in Figure 4A. Apart from no activation in the primary motor areas, the other activated areas closely matched those reported in the prior physical training study that the current study was based on (Wiestler and Diedrichsen, 2013). The activated bilateral frontoparietal regions largely correspond to the action observation network identified in previous studies (Caspers et al., 2010; Cross et al., 2009; Molenburghs et al., 2012; Kirsch & Cross, 2015). Brain activity maps of Trained and Untrained conditions pre- and post-training are visualised in Figure 4B.

**Figure 4.**
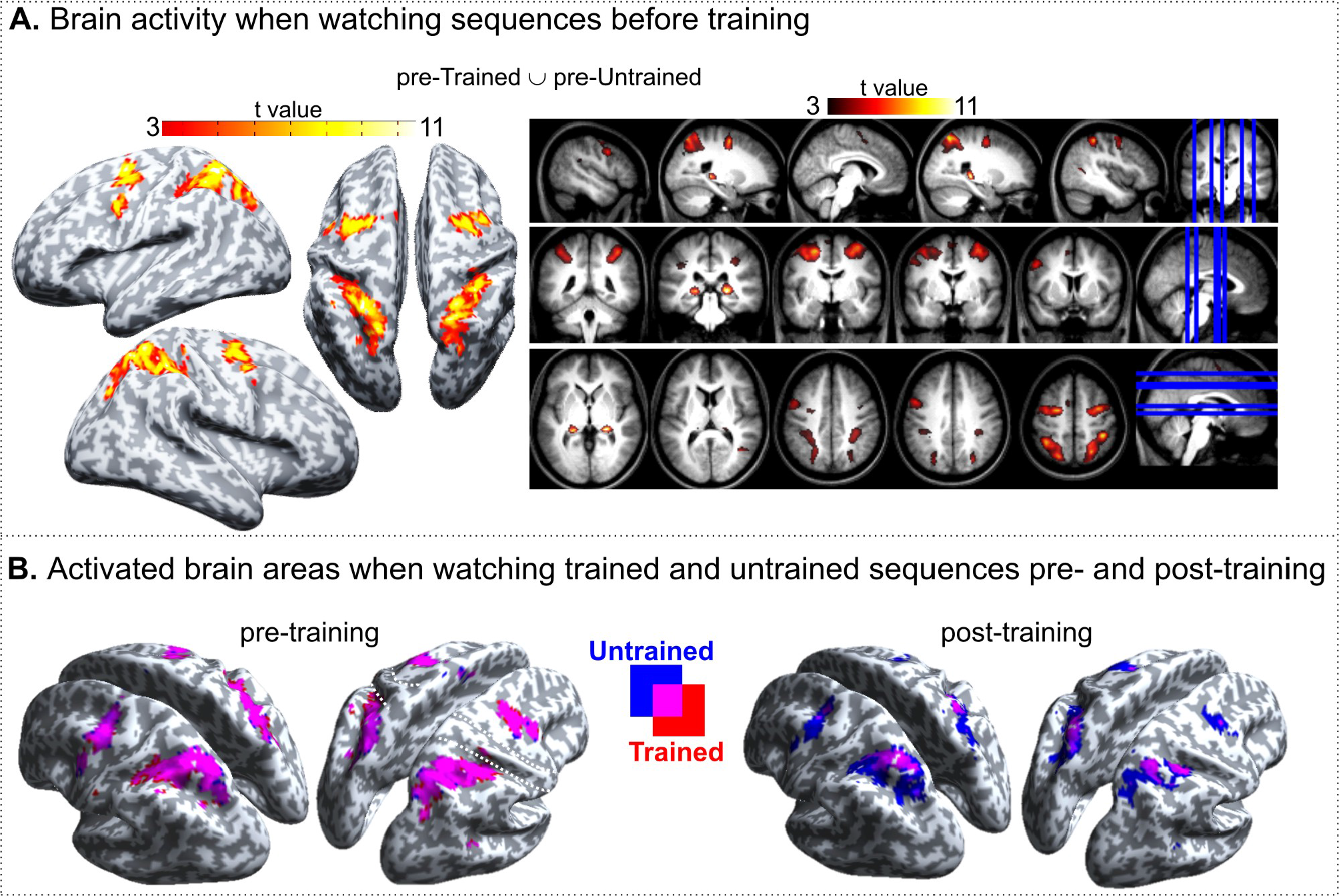
Univariate results. **A.** Activated brain regions when watching sequences before the training (pre-Trained ⋃ pre-Untrained > implicit baseline). Statistical maps are overlaid on inflated standard MNI cortical surface (SPM12) and a group-average T1-weighted image in MNI template space. Maps are thresholded at a single voxel level p < 0.001 (uncorrected, k = 10), showing only clusters with cluster FWE-corrected significance at p < 0.05. **B.** Brain activity maps of Trained (red) and Untrained (blue) conditions pre* and post-training. Maps are thresholded at a single voxel level p < 0.001 (uncorrected), k = 10.

**Table 1.**
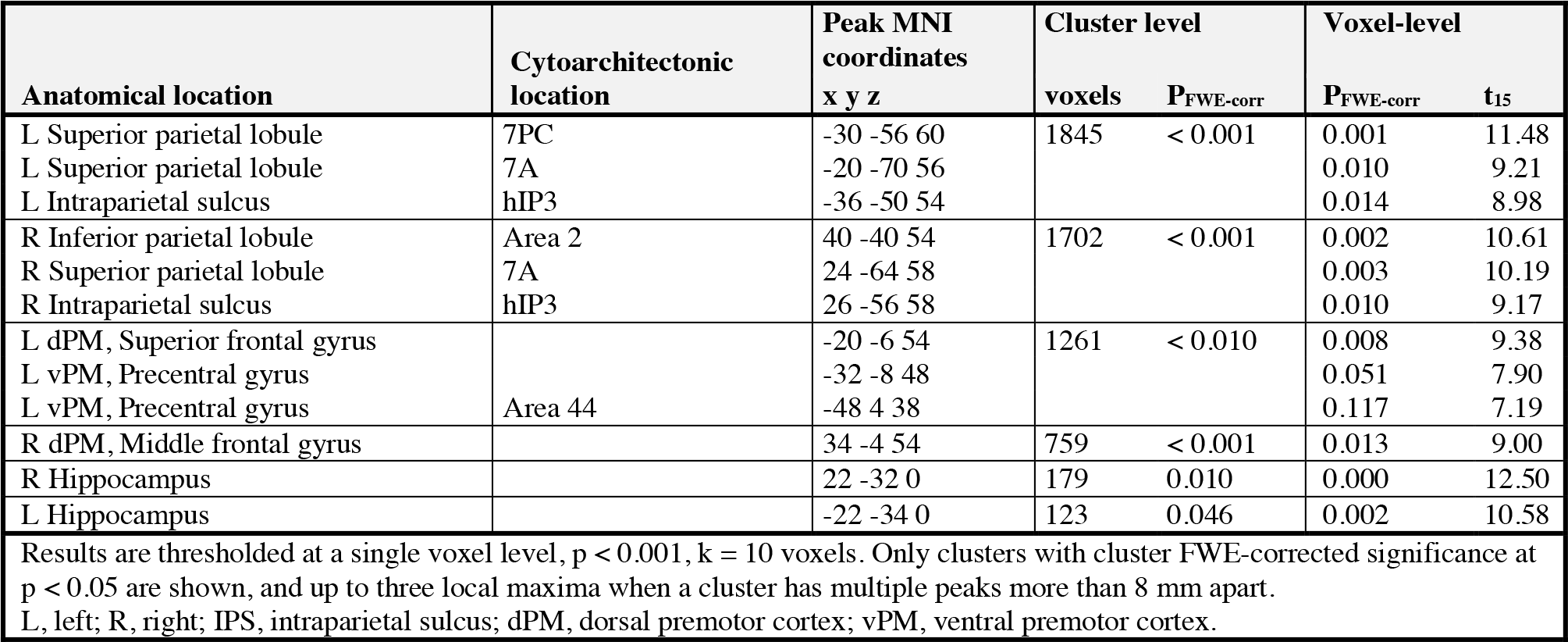
Activated brain regions when watching sequences before the training (pre-Trained ⋃ preUntrained > implicit baseline).

##### Brain regions sensitive to observational training

The post-Untrained > post-Trained contrast revealed clusters in the right superior parietal lobule (extending across right precuneus and left superior and inferior parietal lobules), bilateral dorsal premotor cortices, and left ventral premotor cortex (Table 2 and Figure 5A). After the four days of observational training, therefore, these brain regions showed decreased brain activity when watching trained compared to untrained sequences, which is consistent with prior physical training effects using the same sequences (Wiestler and Diedrichsen, 2013) and observational learning studies using similar sequence learning paradigms (Sakreida et al., 2018). No regions with higher activity for trained compared to untrained were found (as in Wiestler and Diedrichsen, 2013).

**Figure 5.**
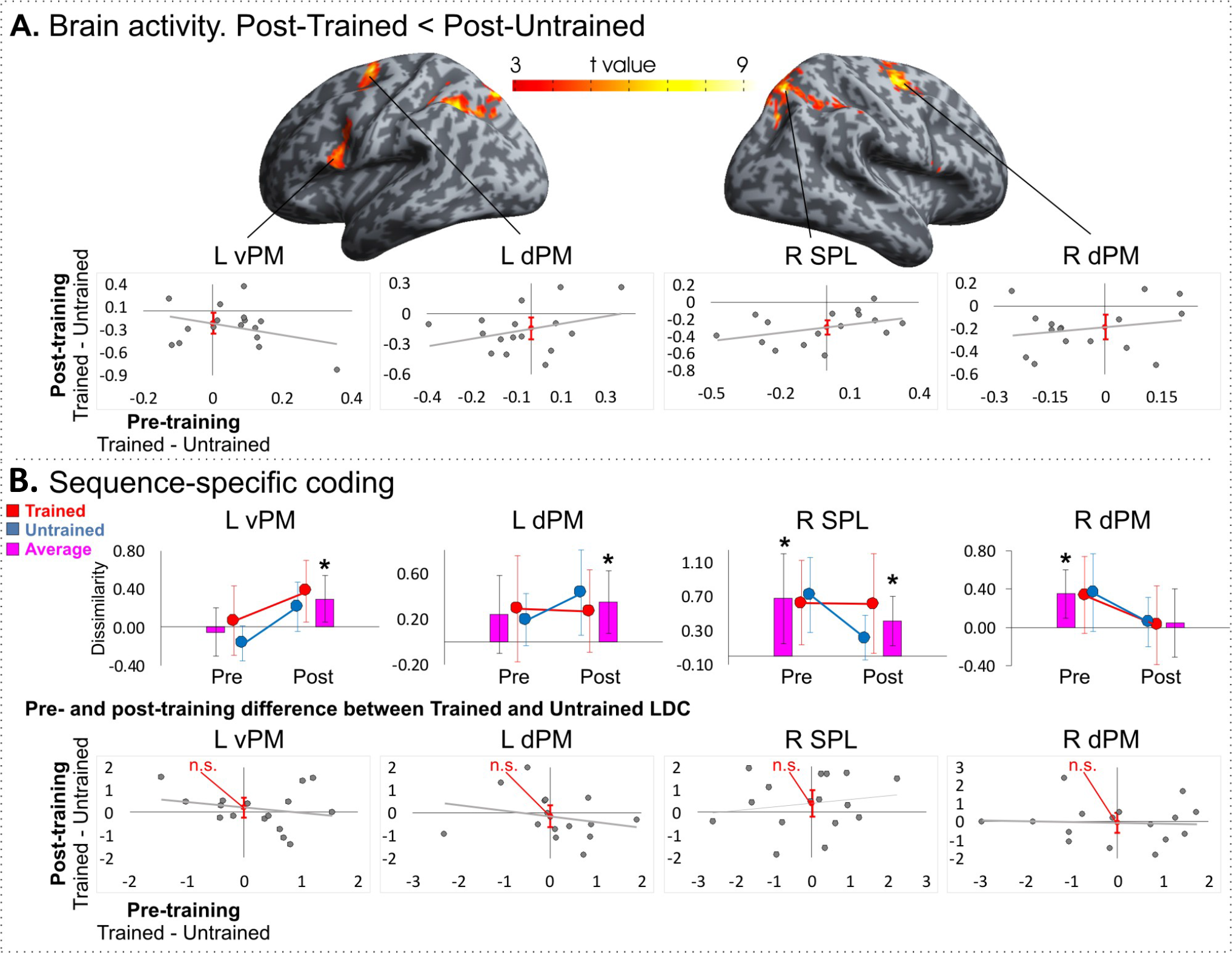
Univariate and multi-voxel pattern analysis region-of-interest (ROI) results. **A.** Univariate results of post-training difference between Trained and Untrained conditions, corrected for pre-training difference. Maps are thresholded at a single voxel level p < 0.001 (uncorrected), k = 10, showing only clusters with cluster FWE-corrected significance at p < 0.05. Plots illustrate pre- and post-training difference in beta weights between Trained and Untrained conditions in the four significant regions selected for further ROI analyses. Error bars represent 95% CI of the intercept. **B.** Top panel: MVPA results of sequence-specific coding pre- and post-training in the four ROIs, showing dissimilarity estimate (average LDC value) between the sequences within the Trained and Untrained conditions and across both conditions on average. Error bars represent within-subject 95% CI; * p < 0.05. Bottom panel: Pre- and post-training difference between Trained and Untrained LDC. Error bars represent 95% CI of the intercept; n.s. – non-significant. L, left; R, right; vPM, ventral premotor cortex; dPM, dorsal premotor cortex; SPL; superior parietal lobule.

**Table 2.** Brain regions showing lower activity for trained compared to untrained sequences post training.

| Anatomical location | Cytoarchitectonic location | Peak MNI coordinates | Cluster level |  | Voxel-level |  |
| --- | --- | --- | --- | --- | --- | --- |
|  |  | x y z | voxels | P<sub>FWE-corr</sub> | P<sub>FWE-corr</sub> | t<sub>14</sub> |
| **R Superior parietal lobule** | **7A** | **22 -68 56** | **1710** | **< 0.001** | **0.007** | **9.43** |
| R Precuneus |  | 10 -58 48 |  |  | 0.068 | 7.86 |
| L Intraparietal sulcus | hIP3 | -28 -50 40 |  |  | 0.210 | 7.16 |
| **R dPM, Superior frontal gyrus** |  | **30 -4 58** | **610** | **< 0.001** | **0.049** | **8.07** |
| R dPM, Precentral gyrus |  | 28 -6 50 |  |  | 0.066 | 7.88 |
| R dPM, Posterior-medial frontal cortex |  | 16 -4 62 |  |  | 0.979 | 5.09 |
| **L vPM, Inferior frontal gyrus (opercularis)** | **Area 44** | **-44 2 24** | **372** | **< 0.001** | **0.708** | **5.94** |
| L vPM, Inferior frontal gyrus (opercularis) | Area 44 | -56 8 10 |  |  | 0.891 | 5.50 |
| L vPM, Precentral gyrus | Area 44 | -50 6 20 |  |  | 0.958 | 5.24 |
| **L dPM, Superior frontal gyrus** |  | **-24 -4 60** | **321** | **< 0.001** | **0.044** | **8.14** |
| L dPM, Middle frontal gyrus |  | -24 -6 50 |  |  | 0.814 | 5.71 |
| L dPM, Middle frontal gyrus |  | -12 -4 58 |  |  | 0.994 | 4.88 |
| Results are thresholded at a single voxel level, $p < 0.001$, $k = 10$ voxels. Only clusters with cluster FWE-corrected significance at $p < 0.05$ are shown, and up to three local maxima when a cluster has multiple peaks more than 8 mm apart. Highlighted are the highest peaks within each cluster which were selected for ROI analyses.<br>L, left; R, right; IPS, intraparietal sulcus; dPM, dorsal premotor cortex; vPM, ventral premotor cortex.<br>The opposite contrast (post-Trained > post-Untrained) did not result in any significant clusters. |  |  |  |  |  |  |

We hypothesised that brain regions that show decreased activity following training would also show distinctive patterns of activity for different sequences in general, as well as for trained compared to untrained sequences (Figure 1E). To investigate this hypothesis, we performed a MVPA on the four ROIs that showed a reduced BOLD response for trained compared to untrained sequences. In addition, we performed an exploratory MVPA using a searchlight approach across the whole brain.

#### Multi-voxel pattern analysis results: sequence-specific representations of observed actions

LDC analyses were used to test whether brain regions hold sequence-specific information following the observation of action sequences and whether the coding of such information is more distinct for trained compared to untrained sequences. The average dissimilarity (LDC value) of activity patterns between the four sequences within each condition was used as a measure of sequence-specific representations.

##### Multi-voxel pattern analysis - ROI approach

We evaluated four ROIs that were sensitive to observational practice (Table 2, Figure 5A): right superior parietal lobule, right dorsal premotor cortex, left ventral premotor cortex, and left dorsal premotor cortex. Each ROI contained an average of 325 voxels (SD = 48.83). On average across Trained and Untrained conditions post-training, sequence-specific activity patterns were found in the right superior parietal lobule, left ventral premotor cortex, and left dorsal premotor cortex (Figure 5B, Table 3). More specifically, sequence-specific activity patterns were found in the right superior parietal lobule, both at pre- and post-training; right dorsal premotor cortex only at pretraining; and left ventral premotor and left dorsal premotor cortices only at post-training. The effect sizes were medium to large in magnitude according to Cohen’s benchmark criteria (Cohen, 1992), as they ranged from Cohen d_z_ = 0.65 to 0.77 (Table 3). These results show that parts of frontoparietal cortex that show sensitivity to observational learning, as measured by changes in average activity, also show distinctive patterns of activity as a function of observed keypress sequences.

**Table 3.** Sequence-specific coding in regions of interest.

| ROI | Mean LDC, One-sample, one-tailed t-test |  |  |  |  | Post-Trained versus Post-Untrained |
| --- | --- | --- | --- | --- | --- | --- |
| R SPL | Pre: 0.68 [0.24, 1.11] | $t_{15} = 2.7$ | $p = 0.008$ | $d_z = 0.68$ | | $B_0 = 0.41 [-0.22, 1.05]$, n.s., $d = 0.35$ |
| | Post: 0.42 [0.18, 0.65] | $t_{15} = 3.08$ | $p = 0.004$ | $d_z = 0.77$ | | |
| R dPM | Pre: 0.35 [0.14, 0.56] | $t_{15} = 2.91$ | $p = 0.005$ | $d_z = 0.73$ | | $B_0 = -0.04 [-0.64, 0.57]$, n.s., $d = 0.03$ |
| | Post: 0.04 [-0.25, 0.33] | n.s. | | $d_z = 0.06$ | | |
| L vPM | Pre: -0.05 [-0.26, 0.16] | n.s. | | $d_z = 0.11$ | | $B_0 = 0.22 [-0.26, 0.70]$, n.s., $d = 0.25$ |
| | Post: 0.29 [0.10, 0.49] | $t_{15} = 2.59$ | $p = 0.01$ | $d_z = 0.65$ | | |
| L dPM | Pre: 0.24 [-0.04, 0.52] | n.s. | | $d_z = 0.38$ | | $B_0 = -0.14 [-0.66, 0.39]$, n.s., $d = 0.14$ |
| | Post: 0.35 [0.12, 0.58] | $t_{15} = 2.69$ | $p = 0.008$ | $d_z = 0.67$ | | |
| LDC, Linear discriminant contrast; L, left; R, right; SPL, Superior parietal lobule; dPM, dorsal premotor cortex; vPM, ventral premotor cortex; n.s., non-significant. |  |  |  |  |  |  |

In the same ROIs, there was only suggestive evidence that sequence-specific representational dissimilarity was different between Trained and Untrained sequences at the post-test (Table 3, Figure 5B). In two ROIs, there was a trend towards sequence-specific representations in frontoparietal cortex showing training-specific effects. In these two ROIs, the training-specific effects at the post-training scan (trained > untrained) were small to medium in size (Cohen’s d = 0.25 for left ventral premotor cortex and 0.35 for superior parietal cortex). However, none of the ROIs showed a significant effect of training. Therefore, four days of observational training produced relatively weak evidence that regions of frontoparietal cortex develop distinctive sequence-specific patterns of activity when observing trained compared to untrained sequences.

##### Multi-voxel pattern analysis - searchlight approach

Whole-brain exploratory surface-based searchlight analysis revealed pre-training (averaged across pre-Trained and pre-Untrained conditions) sequence-specific activity patterns in the right anterior intraparietal sulcus and posterior superior parietal lobule (Table 4; Figure 6A, left panel). In addition, post-training (averaged across post-Trained and post-Untrained conditions), sequence-specific activity patterns were found in bilateral supramarginal gyri, anterior intraparietal sulci (homologous to macaque AIP; Culham et al., 2006), left anterior superior parietal lobule, left primary motor and somatosensory cortices, and right parietal operculum (Table 4; Figure 6A, right panel).

**Figure 6.**
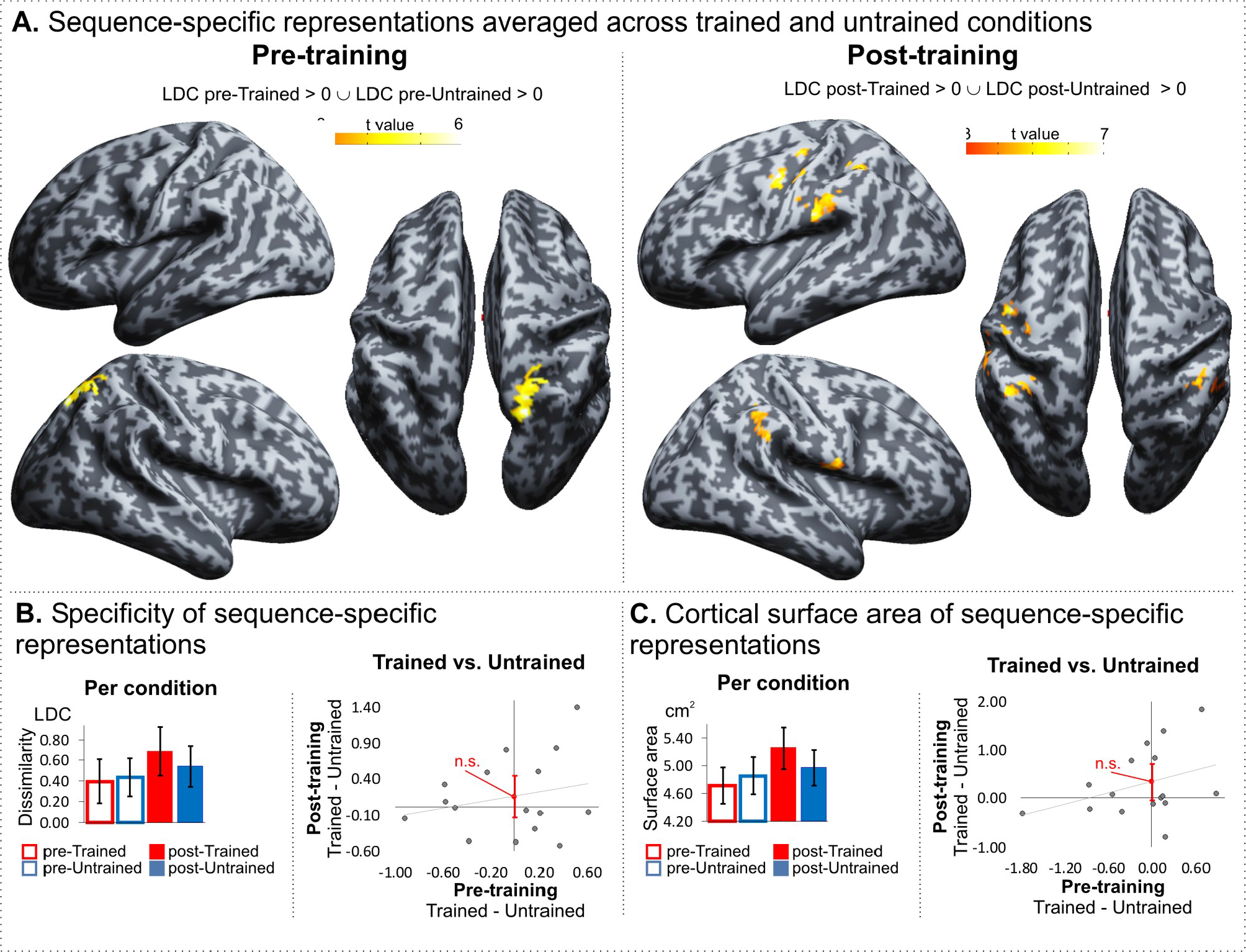
MVPA searchlight results. **A.** Pre-training and post-training sequence-specific representations. Maps are thresholded at a single voxel level p < 0.001 (uncorrected), k = 10, showing only clusters with cluster FWE-corrected significance at p < 0.05. **B.** and **C.** Specificity (the average LDC measure) of sequence-specific representations and the cortical surface area coding sequence-specific representations averaged over all involved cortical regions per condition (left; Error bars represent within-subject 95% CI; * p < 0.05) and pre- and post-training difference (right; Error bars represent 95% CI of the intercept; n.s.: non-significant at p < 0.05).

**Table 4.** Brain regions showing sequence-specific coding for Trained ⋃ Untrained conditions pre- and post-training.

|  |  | Peak MNI coordinates | Cluster level |  | Voxel-level |  |  |
| --- | --- | --- | --- | --- | --- | --- | --- |
| Anatomical location | Cytoarchitectonic location | x y z | voxels | P<sub>FWE-corr</sub> | P<sub>FWE-corr</sub> | t<sub>15</sub> | Average LDC |
| Pre-training |  |  |  |  |  |  |  |
| R Intraparietal sulcus | hIP3 | 22 -62 58 | 453 | < 0.001 | 0.543 | 5.88 | 0.95 |
| R Superior parietal lobule | 7A | 20 -68 50 |  |  | 0.590 | 5.79 | 1.02 |
| R Superior parietal lobule |  | 20 -56 48 |  |  | 0.914 | 5.04 | 0.52 |
| Post-training |  |  |  |  |  |  |  |
| L Supramarginal gyrus | PFop | -56 -26 22 | 269 | 0.001 | 0.377 | 6.32 | 0.82 |
| L Supramarginal gyrus | PFt | -56 -24 32 |  |  | 0.949 | 4.96 | 0.74 |
| L Supramarginal gyrus | PFt | -66 -26 38 |  |  | 0.995 | 4.53 | 0.30 |
| L M1, Precentral gyrus | 4a | -50 -10 42 | 157 | 0.020 | 0.170 | 7.04 | 0.77 |
| L M1, Postcentral gyrus | 4p | -42 -8 34 |  |  | 0.849 | 5.29 | 0.32 |
| L S1, Postcentral gyrus | 3b | -46 -16 48 |  |  | 0.994 | 4.57 | 0.88 |
| R Intraparietal sulcus | hIP2 | 48 -38 42 | 145 | 0.029 | 0.971 | 4.83 | 0.96 |
| R Supramarginal gyrus | PF | 58 -40 30 |  |  | 0.997 | 4.46 | 0.74 |
| R Inferior parietal lobule | Area 2 | 48 -36 52 |  |  | 1.000 | 4.24 | 0.71 |
| L Intraparietal sulcus | hIP2 | -46 -48 54 | 143 | 0.030 | 0.907 | 5.12 | 0.92 |
| L Superior parietal lobule | 5L | -32 -42 46 |  |  | 0.970 | 4.48 | 0.55 |
| R Parietal operculum | OP4 | 58 -8 12 | 134 | 0.039 | 0.874 | 5.22 | 0.70 |
| Results thresholded at a single voxel level, p < 0.001, k = 10 voxels. Only clusters with cluster FWE-corrected significance at p < 0.05 are shown, and up to three local maxima when a cluster has multiple peaks more than 8 mm apart.<br>L, left; R, right; M1, Primary motor cortex; S1, Primary somatosensory cortex; S2, Secondary somatosensory cortex; IPL, Inferior parietal lobule; IPS, Intraparietal sulcus; SPL, Superior parietal lobule. |  |  |  |  |  |  |  |

Similar to the ROI analyses, there was only suggestive evidence that sequence-specific activity patterns become more distinct when watching Trained compared to Untrained sequences following observational training. For exampe, at a cluster FWE-corrected threshold of p < 0.05, no brain regions showed sequence-specific and training-specific patterns of activity. In addition, when comparing the sequence-specific activity patterns globally, averaging over all involved cortical regions, there was only suggestive evidence for more distinct and more widespread sequence-specific coding following practice. Specifically, the average LDC measure of the post-Trained sequences was higher than of the post-Untrained sequences, however the difference was not significant, t_14_ = 1.128, p = 0.278, d_z_ = 0.28, B_0_ = 0.155 [-0.139, 0.449] (Figure 6B). Similarly, the average cortical surface area coding sequence-specific representations of the post-Trained sequences was larger than of the post-Untrained, but the difference was not significant, t_14_ = 1.935, p = 0.073, d_z_ = 0.48, B_0_ = 0.34 cm^2^ [-0.035, 0.715] (Figure 6C).

## Discussion

The neural changes that underpin how visual signals are mapped onto motor circuits when we learn by observation have remained largely unclear. Here we show that observed action sequences are modelled by distinct patterns of activity in frontoparietal cortex and that such representations largely generalise to very similar, but untrained, sequences. These findings advance our understanding of what is modelled during observational learning (sequence-specific information), as well as how it is modelled (reorganisation of frontoparietal cortex is similar manner to that of physical practice). Thus, on a more fine-grained neural level than demonstrated previously, we show the representational structure of how frontoparietal cortex maps visual information onto motor circuits to order to enhance motor performance.

### Sequence-specific activation patterns in frontoparietal brain regions during the observation of action sequences

Prior work has shown that physically practicing keypress sequences leads to reduced engagement and patterns of activity that are sequence-specific in frontoparietal cortex (Wiestler & Diedrichsen, 2013). Here, we show that observation of action sequences also leads to a similar functional reorganisation of frontoparietal cortex. Right SPL, Left PMd and PMv showed a reduction in engagement after visual training of these sequences (Vogt et al., 2007; Higuchi et al., 2012; Sakreida et al., 2018), as well as sequence-specific patterns of activity. The results show close correspondence to prior work on physical practice (Wiestler & Diedrichsen, 2013), by demonstrating that similar regions that distinguish between physically practiced sequences also show sequence-specific patterns when sequences are trained via observation. Moreover, the searchlight analysis showed that premotor and parietal cortices, rather than primary motor cortex, showed sequence-specific representations. As such, similar levels of the motor system hierarchy (Abrahamse et al., 2013; Diedrichsen & Kornysheva, 2015) appear to be modified following physical and observational exposure to action sequences. Thus, we show that patterns of activity in frontoparietal cortices represent action sequences in a similar manner whether the action sequences are physically performed or observed.

The results update our understanding of the role of frontoparietal cortex in shared representations between action and perception in general (Gentsch et al., 2016), as well as our understanding of the features modelled during observational learning (Blandin et al., 1999; Hodges et al., 2007; Boutin et al., 2010). Prior work has shown that action observation and performance share cognitive and neural mechanisms (Gentsch et al., 2016; Giese & Rizzolatti, 2015; Prinz, 1997), which span different levels of the motor hierarchy (e.g., intentions, goals, motor commands; Grafton & Hamilton, 2007). In the present study, sequences were similar to each other at all levels of the motor hierarchy (intentions, goals, motor commands), and differed only in the sequential order of keypresses. Despite the close similarity between the individual actions, we found sequence-specific representations in right SPL, left PMd and PMv. This result deepens understanding of what is shared between perception and production of action (de Vignemont & Haggard, 2008). Rather than observed sequences being represented on a coarser scale (5 keypresses in any order, for example), they are discriminable at an individual sequence level in frontoparietal cortex. This finding thus demonstrates how motor circuit involvement in perception of action sequences has high fidelity to the physical performance.

### Neural plasticity following observational practice

Behavioural data show that observational training leads to faster initiation and movement times for trained compared to untrained sequences and decreases in neural activity within frontoparietal cortex, which mirrors results from physically practicing identical sequences (Wiestler & Diedrichsen, 2013). Therefore, in terms of averaged activity, similar neural efficiency or redundancy gains were seen following observational practice as physical practice (Higuchi et al., 2017; Wiestler & Diedrichsen, 2013). In addition, we show similar evidence of neural generalisation following training: sequence-specific representations were measurable when observing trained and untrained sequences after four days of training, which replicates prior physical training effects (Wiestler & Diedrichsen, 2013) and is consistent with our behavioural data. These data show clear evidence of generalisation of learning from trained to untrained sequences. Given that these sequences were visual and motorically very similar to each other and many of the trained sequences had similar finger transitions to the untrained sequences (Wiestler & Diedrichsen, 2013), it was expected that generalisation would occur. Indeed, it is likely that learned transitions are “chunked” during learning and therefore benefit performance when those transitions between key-presses are present in the untrained sequences (Wymbs et al., 2012).

Wiestler and Diedrichson (2013) also showed that physical practice leads to more distinct sequence-specific representations for trained compared to untrained sequences in frontoparietal cortex. The current study only shows suggestive evidence that following observational learning sequence-specific representations in frontoparietal cortex are more distinctive for trained compared to untrained sequences. For example, in our ROI approach, training-specific effects of MVPA were relatively small (Cohen’s d 0.25 and 0.35) and did not reach a pre-defined statistical threshold of p <0.05. In addition, although evidence in support of training-related differences in representational distinctiveness was relatively weak, the whole-brain searchlight approach also suggested that frontoparietal cortex develops training-specific and sequence-specific representations, which covers a greater proportion of cortex following training. Although more robust than what we report here, it is worth noting that the effects of physical practice in prior work were also rather subtle (Wiestler and Diedrichsen, 2013). In a global analysis that averaged activity in frontal and parietal areas, the effect of physical training corresponded to a 4% increase. Further, in a map-wise analysis, only left SMA / pre-SMA showed a reliable effect for trained compared to untrained sequences. Given that the behavioural effect of observational learning is smaller than physical learning, it is possible that observational learning results in more distinctive sequence-specific patterns of activity but the effect sizes are smaller than physical practice and therefore harder to detect. Given the similarity in behavioural training effects between physical and observational learning of sequences, as well as the similarity in magnitude-based measures of neural activity in frontoparietal cortex, we suggest that this interpretation is likely. Alternatively, it is possible that observational learning does not lead to modified patterns of activity that are sequence and training specific in a manner similar to physical learning. Only future research will be able to confirm or deny these possibilities.

Together, these findings point towards a more general insight into the functional reorganisation of frontoparietal cortex following observational learning. If only univariate results are considered, then reduced engagement of frontoparietal cortex is consistent with greater efficiency in neural function: reduced and less widespread neural engagement is associated with improved physical performance (Steele and Penhune, 2010). However, by unpacking the representational structure of frontoparietal cortex in a sequence-specific manner, we are able to show that frontoparietal cortex develops a richer and more widespread representation of observed action sequences, which largely generalises to untrained sequences. Previous research based on averaging activity across voxels has fuelled much debate about the relative contribution of increased or decreased engagement of the motor system in learning (Dayan & Cohen, 2011; Gardner, Aglinskas & Cross, 2017; Steele and Penhune, 2010). Extending this work, here we emphasise that unlocking the code that is hidden within averaged activity can provide an altogether different understanding of brain organisation (Kriegeskorte, 2008; Norman, 2006). Moreover, the results highlight the value of using representational similarity analyses in the context of learning to understand plasticity, which few studies have focussed upon to date (Kriegeskorte & Kievit, 2013).

## Limitations

In the present study, all eight sequences (four to-be-trained and four untrained) were physically performed before the four days of observational training. Thus, the post-training performance improvement, at least partly, could be driven by the consolidation of physical performance (Censor et al., 2012). While some contribution of physical practice is possible, there was considerably more observational practice (100s of observations per sequence vs. 5 executions). Moreover, trained and to-be-untrained sequences were all physically practiced before the first scan, so any comparisons between trained and untrained sequences were matched for physical practice. For these reasons, we do not think that physical practice had a substantial influence on training-specific effects.

Differences between the current results and those obtained previously from physical practice may result from different dose-response relationships. Although the behavioural and univariate effects of observational training were quite large, the potency of observational practice is likely to be less than physical practice. Therefore, if we had provided sufficient training through observational practice to match the behavioural training gains following physical practice, an even closer set of results may emerge between observational and physical training. Further, we also acknowledge that participants in the current study were not told to intentionally learn the observed action sequences. Instead, participants were told to detect errors. As such, it is possible that the training effects would be larger if participants were given a clear intention to learn. Nonetheless, it remains clear that unintentional learning leads to the type of cognitive and neural re-organisation, which has been outlined in the present paper. Future work that investigates the effect of intentionality in learning using representational similarity analyses would be of interest.

## Acknowledgments

This work was supported by the Ministry of Defence of the United Kingdom Defence Science and Technology Laboratory [grant number DSTLX-1000083177 to ESC and RR], the Economic and Social Research Council [grant numbers ES/K001884/1 to RR and ES/K001892/1 to ESC], and a Marie Curie Actions/FP7 [CIG11-2012-322256 to ESC].

